# Prostaglandins as Candidate Ligands for a Per-ARNT-Sim (PAS) Domain of Steroid Receptor Coactivator 1 (SRC1)

**DOI:** 10.1101/2023.07.13.548854

**Authors:** Nicolas Daffern, Kade Kelley, José A. Villegas, Ishwar Radhakrishnan

## Abstract

Steroid receptor coactivators (SRCs) comprise a family of three paralogous proteins commonly recruited by eukaryotic transcription factors. Each SRC harbors two tandem Per-ARNT-Sim (PAS) domains that are broadly distributed that bind small molecules and regulate interactions. Using computational docking, solution NMR, mass spectrometry, and molecular dynamics simulations, we show that the SRC1 PAS-B domain can bind to certain prostaglandins (PGs) either non-covalently to a surface that overlaps with the site used to engage transcription factors or covalently to a single, specific, conserved cysteine residue next to a solvent accessible hydrophobic pocket. This pocket is in proximity to the canonical transcription factor binding site, but on the opposite side of the domain, suggesting a potential mode of regulating transcriptional activator-coactivator interactions.

## Introduction

Transcriptional coactivators occupy an important niche in promoting gene transcription in eukaryotes. They lack the sequence-specific DNA-binding activity that characterizes transcriptional activators, but commonly serve two critical functions: (i) as molecular adaptors bridging promoter- and enhancer-bound activators with the transcription machinery and (ii) as agents that serve to overcome the repressive effects of chromatin [1, 2]. In the latter instance, coactivators either harbor chromatin-modifying or chromatin-remodeling activities themselves or recruit others with these activities to produce localized perturbations to chromatin structure and dynamics. The molecular functions of coactivators are diverse and go beyond these well documented roles.

The p160 family of steroid receptor coactivators (SRCs, also known as nuclear receptor coactivators or NCoAs) comprising SRC1, SRC2, and SRC3 was originally identified to serve as coactivators of steroid receptors, a family of 6 nuclear receptors (NRs) that bind and respond to steroid hormones [3, 4]. SRCs were subsequently found to serve as coactivators for many of the 42 other human NRs as well as other transcription factors and, as such, constitute one of the most important coactivator families in higher eukaryotes. The SRC proteins, spanning ∼1400 residues in length, are predicted to be largely unstructured but share a common domain architecture. The central region of these proteins harbor 3 highly conserved LxxLL sequence motifs that serve as recruitment sites for NR ligand-binding domains (LBDs), while two separate conserved segments C-terminal to this region serve to engage with the CBP/p300 and the CARM1/PRMT1 coactivators that each harbor distinct chromatin-modifying activities.

The SRC family belongs more broadly to the bHLH-PAS (basic helix-loop-helix Per-ARNT-Sim) family of transcription factors that is dominated by sequence-specific DNA-binding factors as well as a few coactivators [5, 6]. The bHLH-PAS proteins share the same domain architecture at the N-terminus comprising a bHLH DNA-binding/dimerization domain and two PAS domains (designated PAS-A and PAS-B) that also serve as sites for homo-and hetero-dimerization or engagement with other transcription factors and coactivators. Little is known about the function of these domains within the SRC family, although the SRC1 PAS-B domain serves as the coactivator recruitment site for the transactivation domains (TADs) of the STAT6 and Nurr1 transcription factors [7-10].

PAS domains are ubiquitous in biology, and while most of them engage in protein-protein interactions, a subset binds to small molecule ligands that function as co-factors to modulate interactions or transduce signals [11-13]. Indeed, the PAS domains of bHLH-PAS family members are known to harbor internal pockets and a few, including the arylhydrocarbon receptor (AHR) and hypoxia inducible factor-2α (HIF-2α), have been shown to bind to small molecule ligands, activating these intracellular receptors and altering the stability of protein-protein complexes [14, 15]. Here, we show that the SRC1 PAS-B domain has conserved pockets and can bind to specific small molecule ligands including certain prostaglandins via diverse mechanisms.

## Materials and Methods

### Protein expression, purification, and NMR sample preparation

^15^N-labeled SRC1 PAS-B and SRC1 PAS-B-Nurr1 AF1^28-51^ fusion proteins were expressed in bacteria and purified exactly as previously described [10]. NMR samples of the purified proteins at 0.2 mM concentration were prepared in NMR compatible buffer containing 20 mM sodium phosphate (pH 6.8), 50 mM NaCl, 1 mM DTT, and 0.5 mM EDTA. A C347S mutant construct was engineered via the QuikChange protocol using the plasmid encoding wild-type SRC1 PAS-B as the template. The mutation was confirmed both by DNA sequencing and mass spectrometry of the purified protein (see below).

### NMR spectroscopy and small molecule titrations

2D ^1^H-^15^N heteronuclear single-quantum coherence (HSQC) NMR spectra were acquired on a Bruker Neo 600 MHz spectrometer equipped with a QCI-F cryoprobe at 25 °C sample temperature for SRC1 PAS-B and 30 °C for the SRC1 PAS-B-Nurr1 AF1^28-51^ fusion. NMR spectra were processed using Bruker TopSpin or NMRpipe [16] and analyzed using NMRFAM-Sparky [17]. All small-molecule compounds used for the titrations were purchased from vendors in highly pure form and used without further purification. Prostaglandins (PGs) were purchased from Cayman Chemical while all the others were purchased from Sigma. The compounds were dissolved in DMSO or NMR buffer at high concentrations (≥25 mM). Background chemical shift perturbations (CSPs) caused by DMSO were subtracted prior to quantitation of ligand-induced shifts. CSPs were calculated as 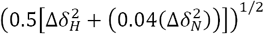, where δ_H_ and δ_N_ are the amide proton and nitrogen chemical shifts, respectively.

### Mass spectrometry

To confirm the identity and purity of wild-type and mutant proteins as well as to assess covalent modification caused by certain PGs, liquid chromatography coupled with electrospray ionization mass spectrometry (LC-ESI-MS) was performed using an Agilent 6545 QTOF instrument. Runs were performed by injecting 7 μl of protein samples in native buffers onto a guard column using a running buffer of 0.1% (v/v) formic acid in H_2_O. Proteins were then eluted with a running buffer of 100% (v/v) acetonitrile for detection. The resulting spectra were deconvoluted and analyzed using Agilent MassHunter BioConfirm software.

### Covalent docking and molecular dynamics (MD) simulations

Calculations were carried out using the Schrodinger suite of programs (Schrödinger, LLC, New York, Release 2020-3). The Covalent Docking module within Glide. was used to dock an atomic model of 15d-PGJ2 (generated using LigPrep and ConfGen) with SRC1 PAS-B (PDB accession: 5Y7W, Chain A, following removal of residues arising from a cloning artifact and processing using Protein Preparation Wizard). The Michael Addition reaction type option was used to generate two protein-bound models, one for each reactive center C9 and C13 ligated to the C347 S^γ^ atom, using Thorough pose prediction mode. The models were then immersed in a 69.7 × 46.7 × 63.8 Å box of SPC water molecules. Counterions as well as Na+ and Cl-ions mimicking an ionic strength of 0.15 M were added. Using the DesmondGPU module in the suite, full-atom NPT ensemble MD simulations were carried out after treatment with the default equilibration protocol. The production run was performed at 300 K for 250 ns using a timestep of 2 femtoseconds. MD trajectories were loaded onto VMD and analyzed using custom scripts. RMSD values from frame 1 for protein were calculated using C^α^ carbons, and for ligands using heavy atoms.

## Results

### SRC1 PAS-B domain harbors a conserved internal pocket

The SRC family exhibits distinct patterns of conservation in the bHLH-PAS region with the bHLH and PAS-A domains being the most strongly conserved in both orthologs and paralogs while the PAS-B domain is highly diversified among paralogs (**Supplementary Figure S1a**). However, among orthologs, the SRC1 PAS-B domain is highly conserved, especially the residues on the surface that engage STAT6 TAD and Nurr1 AF1 (**Figure 1a and 1b**). Analysis of the crystal structures of the domain bound to STAT6 TAD or stapled peptide mimics thereof reveals an internal pocket that is 280 to 380 Å^3^ in volume (**Figure 1b**; [8, 18]). The pocket is adjacent to the STAT6/Nurr1-binding site with several solvent-exposed residues forming the binding site also directly contributing to the pocket. The irregularly shaped pocket is lined by residues that are essentially invariant in SRC1 PAS-B orthologs, suggesting a functionally significant role (two residues located at the mouth of the pocket, although not invariant, are highly conserved). The pocket is lined by the side chains of both hydrophobic and polar residues (**Figure 1b**; **Supplementary Figure S1b**). A portion of the pocket features a network of three ordered water molecules in the crystal structures that make hydrogen bonding interactions with polar groups in the side chains of T264, Q266, and C344 as well as with the backbone carbonyls of T264, A310, and I358.

**Figure 1.**
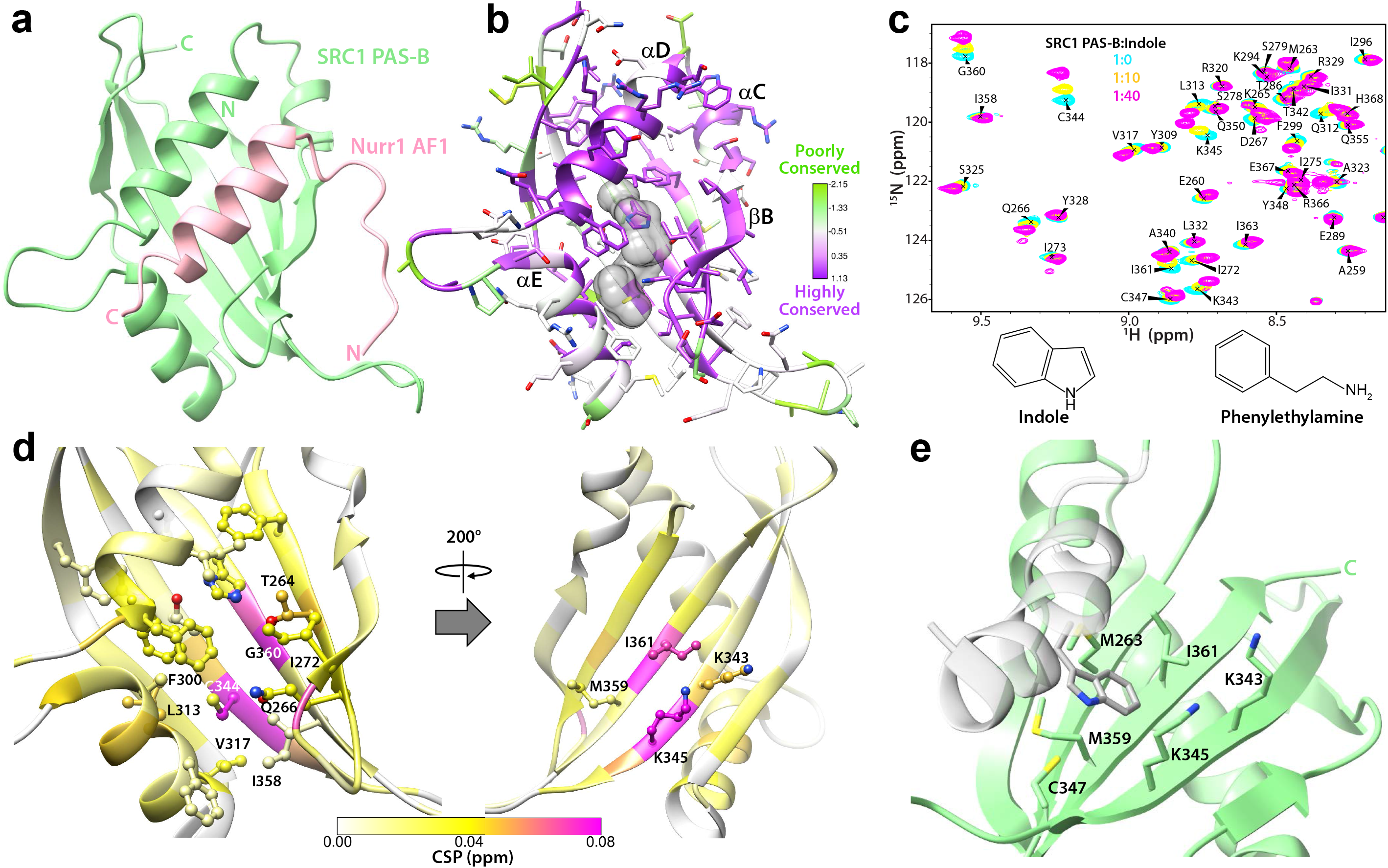
The PAS-B domain harbors conserved internal and external pockets accessible to small ligands. (**a**) An AlphaFold structural model of SRC1 PAS-B (light green) in complex with Nurr1 AF1 (light pink). (**b**) Crystal structure of SRC1 PAS-B in complex with a STAT6 TAD-like stapled peptide (PDB ID:5Y7W) shown with a comparable pose as in panel a; the peptide is not shown for clarity. PAS-B residues are colored by the level of sequence conservation in SRC1 orthologs using Chimera according to the scheme shown in the color key. The internal, irregularly shaped pocket calculated by CASTp is rendered as a semi-transparent, gray colored surface. (**c**) Expanded plots of ^15^N-SRC1 PAS-B ^1^H-^15^N HSQC spectra recorded in the absence and presence of the indicated amounts of indole according to the color key. The bottom panel shows the molecular structures of the ligands that induced significant perturbations in PAS-B spectra. (**d**) Chemical shift perturbations (CSPs) induced by indole at 1:40 protein:ligand ratio mapped on to the PAS-B solution structure (PDB ID: 5NWM); front and back views of the domain are shown. The side chains of residues lining the internal and external (see below) pockets that show some of the strongest CSPs are shown. (**e**) Details of the interaction between the sidechain of a tryptophan residue in the cloning artifact (colored in gray) and the shallow, hydrophobic pocket formed by the side chains of M359, I361, K343, and K345.

### SRC1 PAS-B can specifically bind small molecule compounds

To test whether the pocket was accessible to small molecule ligands, we screened a variety of compounds for binding to ^15^N-PAS-B by solution NMR. Since Nurr1 is a well-established regulator of dopamine biosynthesis, the initial compounds we selected were those that were either precursors, metabolites, or analogs of dopamine (**Supplementary Table 1**). Among this panel of compounds, indole, and to a lesser extent, phenylethylamine caused small but specific chemical shift perturbations (CSPs) in the PAS-B HSQC spectrum (**Figure 1c**). Mapping these perturbations on to the PAS-B structure reveals that residues lining the internal pocket as well as those flanking these residues show the strongest CSPs, suggestive of ligand entering and binding to the pocket (**Figure 1d**) or binding exclusively or additionally to a solvent-accessible, shallow hydrophobic pocket formed by M359, I361, K343, and K345, which like the internal pocket is also highly conserved (**Supplementary Figure S1b**). Notably, in the crystal structure of SRC1 PAS-B bound to a stapled peptide mimic [18], this pocket is occupied by a tryptophan side chain belonging to a cloning artifact at the N-terminus of the domain (**Figure 1e**), demonstrating that this pocket may have a functional role in protein-ligand interactions. Although the PAS-B binding affinities for these compounds are estimated to be in the high millimolar range, making them too weak to be biologically relevant, these results imply that PAS-B can bind small molecule ligands through one of its conserved pockets.

### PAS-B can bind to fatty acids

To identify candidate endogenous ligands for the SRC1 PAS-B domain, we conducted an unbiased *in silico* screen of compounds in the small molecule pathway database (https://smpdb.ca; [19]) by pursuing a flexible docking approach using RosettaLigand [20]. Given the relatively small size of the pocket, we only considered ligands with molecular weight <375 Da. Over 1600 compounds were screened virtually and a handful of the top scoring candidates were evaluated for binding by solution NMR (**Supplementary Table 1**). Although none of these compounds produced any appreciable changes to the HSQC spectrum of ^15^N-PAS-B, visual inspection of the docked poses of ligands with top scores revealed a recurring theme of carboxylate and aliphatic moieties of fatty acids engaging respectively with the side chains of conserved arginine and phenylalanine residues (R311 and F314) in the αE helix (**Figure 1b**).

Prostaglandins are metabolic products of the fatty acid, arachidonic acid, that were shown recently to function as endogenous ligands for Nurr1 LBD. We asked whether PGE1 and PGA1, both of which showed specificity for Nurr1 LBD [21], could bind to the PAS-B domain. Titration with PGE1 produced significant CSPs in the ^15^N-PAS-B HSQC spectrum (**Supplementary Figure S2**); however, the affinity from NMR titrations was estimated to be in the low millimolar range. Incubation of ^15^N-PAS-B with PGA1, which was previously shown to covalently modify Nurr1 LBD [21], resulted in the appearance of new peaks in slow exchange in the HSQC spectrum, consistent with covalent modification but only a small fraction of the protein was modified in this manner even after overnight incubation with the compound. Therefore, we extended our studies to include other PGs related to PGE1 and PGA1 (**Table 1**). Titration with PGB1 produced some of the strongest ligand-induced perturbations in the PAS-B spectrum with the affinity of the interaction estimated to be in the 300 µM range (**Figure 2a**). Structural mapping of the CSPs suggested engagement of the ligand by residues that line the internal pocket (**Figure 2b**). We had shown previously that residues 28-51 within the AF1 transactivation domain of the transcription factor Nurr1 comprised the minimal SRC1 PAS-B binding site [10]. To test whether PAS-B could simultaneously engage Nurr1 AF1^28-51^ and PGB1, titrations were repeated with a ^15^N-PAS-B-Nurr1 AF1^28-51^ fusion construct that we had characterized previously [10]. Although PGB1 induced significant CSPs in the HSQC spectrum of the fusion protein, both the magnitude and pattern of CSPs were different compared to those produced in titrations with PAS-B alone (**Figure 2c; Table 1**). Additionally, PGB1-binding produced CSPs in the Nurr1 AF1 helix with the trajectories of the AF1 peaks during the titration suggesting displacement of the helix and competition between the two binders for overlapping binding sites.

**Table 1.** Chemical Shift Perturbation Statistics for Prostaglandins*

| Protein | Ligand | Mean CSP ± Standard | Mean CSP ± Standard |
| --- | --- | --- | --- |
|  |  | Deviation for All Peaks (ppm) | Deviation for Top 5% of Most Perturbed Peaks (ppm) |
| SRC1<br>PAS-B | PGB1 | 0.052 ± 0.055 | 0.229 ± 0.023 |
| | PGE2 | 0.031 ± 0.031 | 0.129 ± 0.016 |
| | PGA1 | 0.036 ± 0.046 | 0.198 ± 0.029 |
| | PGE1 | 0.032 ± 0.033 | 0.134 ± 0.021 |
| SRC1<br>PAS-B-Nurr1<br>AF1<sup>28-51</sup><br>fusion | PGB1 | 0.026 ± 0.026 | 0.114 ± 0.036 |
| | PGE2 | 0.009 ± 0.007 | 0.031 ± 0.007 |
| | PGA1 | 0.014 ± 0.019 | 0.083 ± 0.027 |
| | 8-iso PGE1 | 0.013 ± 0.010 | 0.041 ± 0.019 |
| | PGD1 | 0.008 ± 0.004 | 0.019 ± 0.002 |
| | 15d-PGJ2 | 0.038 ± 0.064** | 0.270 ± 0.076** |

\*: values are for 1:10 protein:ligand ratio and 0.2 mM protein concentration
\*\* : values are for 1:2 protein:ligand ratio after overnight incubation at 4 °C with 0.2 mM protein

**Figure 2.**
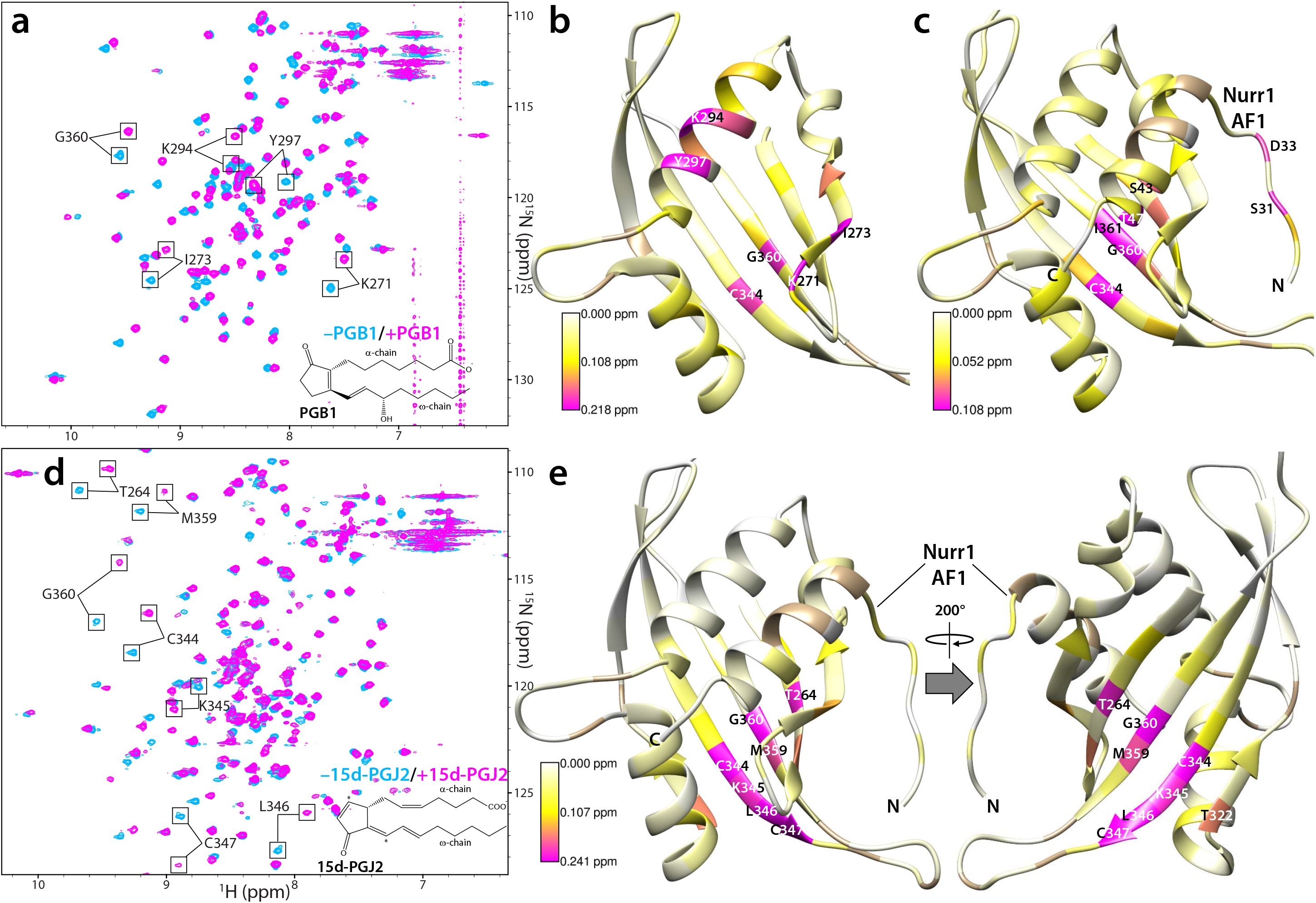
Prostaglandins as candidate ligands for the SRC1 PAS-B domain. (**a**) Expanded plots of ^15^N-SRC1 PAS-B ^1^H-^15^N HSQC spectra recorded in the absence (blue) and presence of 10 equivalents of PGB1 (magenta); the molecular structure of the compound is shown at the bottom. The top 5% of the perturbed peaks are annotated. (**b**) CSPs in the (**b**) ^15^N-PAS-B and (**c**) ^15^N-PAS-B-Nurr1 AF1^28-51^ fusion protein spectra induced by PGB1 at 1:10 protein:ligand ratio mapped on to the PAS-B solution structure (PDB ID: 5NWM) and the AlphaFold model of the PAS-B-AF1^28-51^ complex, respectively. Prolines and line-broadened residues for which CSPs could not be measured are colored in tan. The most strongly perturbed residues are labeled. (**d**) Expanded plots of ^15^N-SRC1 PAS-B-Nurr1 AF1^28-51^ fusion HSQC spectra recorded in the absence (blue) and following overnight incubation at 4 °C with 2 equivalents of 15d-PGJ2 (magenta); the molecular structure of the compound is shown at the bottom. The solution conditions were identical to those described in Figures 1c, 2b and 2c, except the samples lacked DTT. (**e**) CSPs from the spectrum in panel d mapped on to the AlphaFold model of the PAS-B-AF1^28-51^ complex. Front and back views of the domain are shown. Prolines and line-broadened residues for which CSPs could not be measured are colored in tan. The most strongly perturbed residues are labeled.

Incubation of 15-deoxy-Δ^12,14^-PGJ2 (15d-PGJ2) with the ^15^N PAS-B-Nurr1 AF1^28-51^ fusion protein also produced strikingly strong perturbations in the HSQC spectrum (**Figure 2d; Table 1**). Structural mapping of the CSPs suggests that 15d-PGJ2 binds exclusively to the shallow hydrophobic pocket on the opposite side of the domain (**Figure 2e**); the lack of appreciable perturbations in the AF1 helix is indicative of simultaneous binding of both the ligand and Nurr1. Since 15d-PGJ2 can covalently modify cysteine residues via Michael addition ([22]; **Supplementary Figure S3a**) and since there are conserved cysteine residues (C344 and C347) within or in proximity to the two pockets, we asked whether PAS-B could be covalently modified by the ligand. LC-ESI-MS analysis of the NMR sample used to acquire data in Figure 2d shows a major peak with a molecular mass +317 Da higher than the mass expected for the ^15^N-labeled protein (**Supplementary Figure S3a**). This suggests that the protein can be covalently modified by 15d-PGJ2. Although the PAS-B domain contains three cysteine residues (C295, C344, and C347), the covalent modification appears to be restricted to only a single residue, as evidenced by the lack of peaks at higher molecular weights (e.g., +634 Da) in the mass spectrum. While the C295 peak in the PAS-B HSQC spectrum is largely unperturbed, those belonging to C344 and C347 are both strongly affected by the 15d-PGJ2 modification (**Figure 2d & 2e**). To test which of these two residues is preferentially modified by 15d-PGJ2, C347 was mutated to serine in the context of the SRC1 PAS-B construct, and both the purified wild-type and mutant proteins were incubated with the compound overnight. LC-ESI-MS analysis revealed that, although the wild-type protein was singly modified by 15d-PGJ2, as was observed for the ^15^N-labeled PAS-B-Nurr1 AF1 fusion, the C347S mutant was not (**Supplementary Figure S3b**), implying that C347 was the preferred covalent modification site. This result is consistent with the pattern of CSPs induced by the ligand since the peak belonging to C347 is one of the most strongly perturbed (**Figure 2d**). Also strongly perturbed are the peaks belonging to residues that comprise the solvent accessible hydrophobic pocket located adjacent to C347 on the opposite side of the domain (**Figures 1e & 2e**).

### Molecular modeling of 15d-PGJ2 covalently bound to PAS-B

There are two reactive centers in 15d-PGJ2 located at the C9 and C13 positions for Michael addition (**Supplementary Figure S3a**). To establish which of these two centers is likely involved in ligating with C347 S^γ^ and to model how the 15d-PGJ2 moiety is stably accommodated on the surface of PAS-B, 250 ns all-atom molecular dynamics (MD) simulations in explicit solvent, counter-ions, and 150 mM NaCl were performed using the Desmond module in the Schrödinger suite. The atomic root-mean-square deviations (RMSD) for the protein backbone atoms relative to the starting model largely remained below 3 Å in these simulations (**Figure 3a & 3b**). Whereas the PG moiety ligated to C347 via the C13 atom mostly showed RMSDs below 4 Å relative to the starting model, the same moiety ligated via the C9 atom showed RMSDs consistently around 8 Å or above, reflecting the significantly broader diversity of configurational space sampled by the latter compared to the former. This is best exemplified by the salt-bridging interactions engaged by the terminal carboxylate moieties in the two simulations. In the case of 15d-PGJ2 ligated via C13, the carboxylate moiety engages in salt-bridging interactions solely with the ε-ammonium groups of K343 and/or K345 (**Figure 3a**), whereas when ligated via C9, it engages with either K265 or K345 located on disparate parts of the protein (**Figure 3b**). The exclusively aliphatic ω-chain is largely unconstrained in both simulations whereas the aliphatic portion of the α-chain in the case of C13-mediated ligation engages in hydrophobic interactions with the shallow, conserved pocket defined by M359, I361, and the aliphatic segments of K343 and K345 (**Figure 3a and Supplementary Figure S1b**). This feature is consistent with the NMR CSPs (**Figure 2e**), providing support not only for 15d-PGJ2 α-chain targeting the PAS-B surface pocket, but also for a mode of ligation that is mediated by the C13 electrophile.

**Figure 3.**
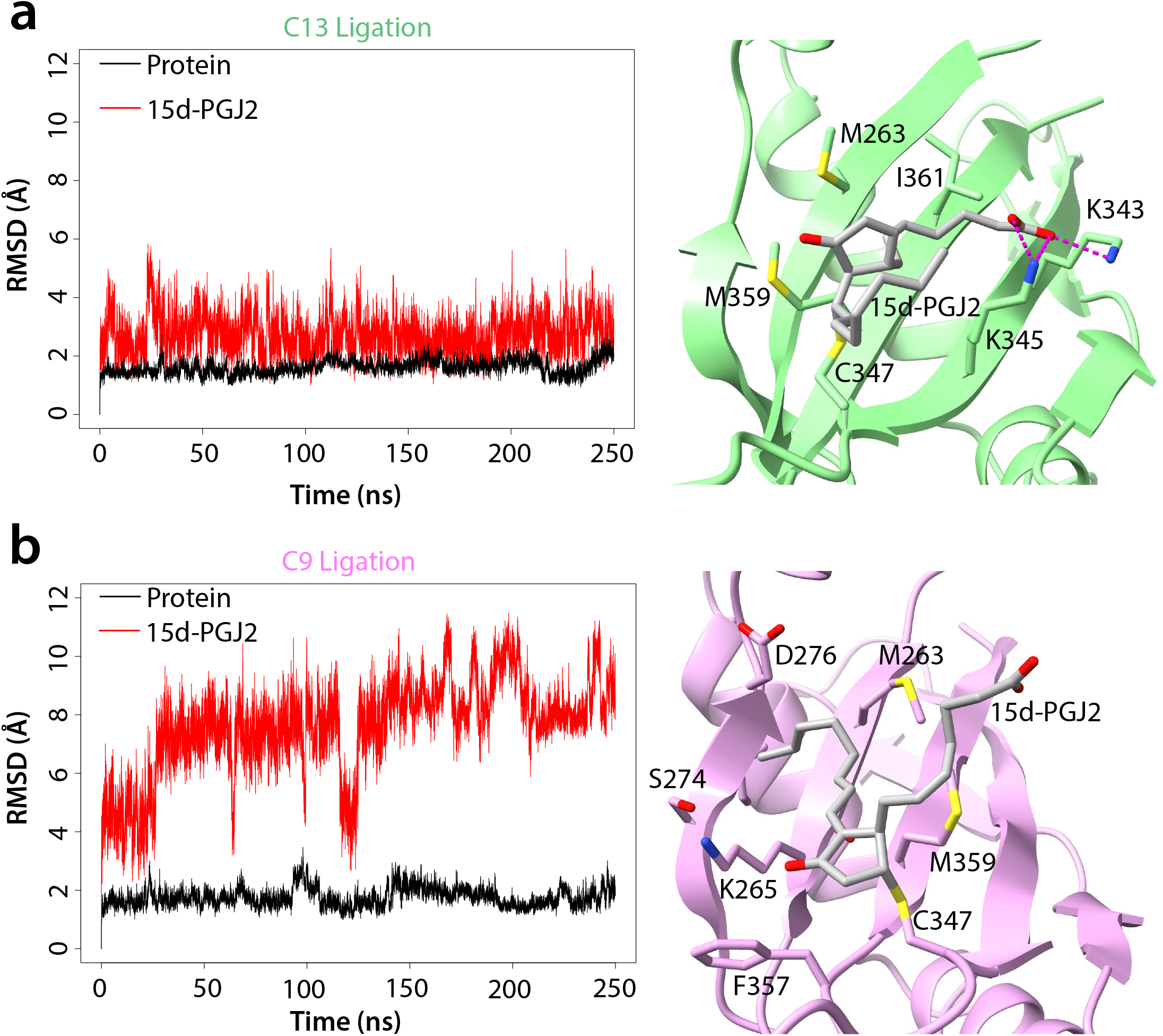
A structural model for 15d-PGJ2 covalently attached via C347 side chain to the SRC1 PAS-B domain. Atomic RMSDs for PAS-B covalently bound via (**a**) C13 and (**b**) C9 atoms to 15d-PGJ2 during 250 ns all-atom MD simulations in explicit solvent, counterions, and salt (*left panels*). The corresponding structural models from the last frame of the MD simulations showing key non-covalent interactions between the PG moiety and the protein (*right panels*).

### Conclusions and biological implications

In conclusion, we have shown that the SRC1 PAS-B domain can bind to small-molecule compounds with an apparent preference for certain fatty acids. Given the proximity of the surface pocket to the site through which the PAS domain commonly engages with diverse transcription factors, it raises the possibility that ligand binding could be a mechanism to modulate the affinity of protein-protein interactions, as has been witnessed for other PAS-domain containing transcription factors [14]. Although our *in vitro* studies identified prostaglandins as potential ligands for SRC1 PAS-B, more comprehensive studies are needed to evaluate the biological impact of these ligands in transcriptional activation by Nurr1 as well as other receptors and transcriptional factors that rely on SRC1 recruitment by engaging with this domain. Also, whereas the interaction with and covalent modification by 15d-PGJ2 appears to be specific, it is possible and indeed likely that other endogenous ligand(s) for the PAS-B domain that target the internal cavity exist, warranting a broader search for such ligands via high-throughput experimental approaches.

Our finding that prostaglandins could be candidate ligands for SRC1 PAS-B is intriguing because PGs play an important role in inducing the inflammatory response as well as in the resolution of this response [23-25]. They are ubiquitously produced in cells and function as autocrine and paracrine mediators to maintain local homeostasis. PGs exert their effects by binding to G protein-coupled receptors and triggering intracellular signaling pathways, but certain PGs can serve as endogenous ligands by binding directly to the LBD of NRs including Nurr1 and PPARγ and stimulating transcriptional activation [21, 22]. PGA1 and 15d-PGJ2 have been implicated in *covalently* binding to Nurr1 and PPARγ, respectively, producing a “permanent” effect on transactivation that in turn is manifested in their anti-inflammatory activities [26]. Indeed, PGA1 and its precursor, PGE1 have been recently shown to protect midbrain dopaminergic neurons from neurotoxins and ameliorate motor deficits in mouse models of Parkinson’s disease [27]. Also, Nurr1 has long been known to protect tyrosine hydroxylase expressing dopaminergic neurons from inflammation-induced cell death [28, 29]. Thus, PGs as candidate potentiators at multiple steps of Nurr1-dependent transactivation, if confirmed, is immensely interesting, not least because of the close ties between these compounds, Nurr1, and inflammation, but also because it would open new therapeutic options for the treatment of Parkinson’s disease.

## Abbreviations

NR: nuclear receptor
LBD: ligand-binding domain
AF1: activation function 1
SRC: steroid receptor coactivator
AHR: arylhydrocarbon receptor
ARNT: arylhydrocarbon receptor nuclear translocator
bHLH: basic helix-loop-helix
PAS: Per-ARNT-Sim
HIF-2α: hypoxia inducible factor-2α
TAD: transactivation domain
HSQC: heteronuclear single-quantum coherence
CSP: chemical shift perturbation
PG: prostaglandins
MD: molecular dynamics
RMSF: root-mean-square fluctuations
15d-PGJ2: 15-deoxy-Δ^12,14^-PGJ2
LC-ESI-MS: electrospray ionization-mass spectrometry

## Acknowledgements

We thank Yongbo Zhang in the IMSERC NMR facility and Saman Shafie in the IMSERC MS facility at Northwestern for technical assistance and advice. This work was supported by a grant from the Sherman Fairchild Foundation to I.R. N.D. was supported by a Molecular Biophysics Training Program traineeship (T32 GM008382). Support for structural biology and biophysics research from the Robert Lurie Comprehensive Cancer Center is gratefully acknowledged.

