## Supplementary Materials for "Prostaglandins as Candidate Ligands for a Per-ARNT-Sim (PAS) Domain of Steroid Receptor Coactivator 1 (SRC1)"

**Supplementary Figure S1.** Sequence Conservation in the N-terminal globular domains of the SRC family and the SRC1 PAS-B domain. (**a**) A Clustal Ω-guided multiple sequence alignment of the N-terminal region encompassing the bHLH, PAS-A, and PAS-B domains in SRC paralogs. Domain boundaries as deduced from the AlphaFold model for SRC1 are shown on top. The sequence identity and similarity metrics resulting from pairwise comparisons of the three proteins for each of the three domains are 61-78% and 78-85%, respectively, for the bHLH domain, 63-77% and 75-84% for the PAS-A domain and 41-52% and 63-75% for the PAS-B domain, making the latter domain the most evolutionarily diverse compared to the other domains. (**b**) A Clustal Ω-guided multiple sequence alignment of the PAS-B domain in SRC1 orthologs. The cartoon above the alignment shows the location of secondary structure elements in the crystal structure. The purple dots at the top of the alignment identify residues comprising the internal pocket while the pink triangles identify residues comprising the external, shallow hydrophobic pocket along with C347.

**Supplementary Figure S2.** SRC1 PAS-B binds weakly to PGE1 that is known to target the Nurr1 LBD. (**a**) $^{1}H$-$^{15}N$ HSQC spectra of 0.2 mM $^{15}N$-PAS-B recorded in the absence and presence of 10 equivalents of PGE1. A stock solution of PGE1 was prepared by dissolving the dry, lyophilized powder in DMSO. NMR data were acquired under the same experimental conditions as described in Figure 1c. The corresponding CSPs for each peak at each titration point were quantified and used to estimate the binding affinity. (**b**) The CSPs at 1:10 protein:ligand ratio were mapped on to the molecular surface of SRC1 PAS-B to deduce the PGE1-binding surface.

**Supplementary Figure S3.** Mass spectra show modification of a specific cysteine residue by 15d-PGJ2. (**a**) Reaction chemistry of Michael addition (*left panel*) and the LC-ESI-MS spectrum of the $^{15}N$-PAS-B-Nurr1 $AF1^{28\text{-}51}$ NMR sample following incubation with 15d-PGJ2 used to record the spectrum in Figure 2d establishing an increase in mass relative to the unmodified protein of 317 Da corresponding to the formation of the 15d-PGJ2 covalent adduct (*right panel*). The two reactive centers at the C9 and C13 carbons in 15d-PGJ2 are labeled; the Michael adduct formed by C13 is shown. (**b**) LC-ESI-MS spectra of wild-type SRC1 PAS-B (*top*) and the C347S mutant (*bottom*) following incubation at 4 °C with 2 equivalents of 15d-PGJ2 overnight.

**Supplementary Table 1.**

**Small-molecule Compounds Tested for Binding to SRC1 PAS-B by NMR***

| Small Molecule | Protein:Ligand Ratios Tested | Protein | Binding (+:Yes/–:No) |
|---|---|---|---|
| 6-mercaptopurine | 1:10 | SRC1 PAS-B | – |
| 5,6-dihydroxyindole** | 1:1 | SRC1 PAS-B | – |
| dimethylsulfoxide | 2.5%, 5% (v/v) | SRC1 PAS-B | + |
| indole | 1:40 | SRC1 PAS-B | + |
| L-tyrosine | 1:1, 1:5 | SRC1 PAS-B | – |
| dopamine | 1:10 | SRC1 PAS-B | – |
| phenylethylamine | 1:10, 1:30 | SRC1 PAS-B | + |
| tyramine | 1:10 | SRC1 PAS-B | – |
| tryptamine | 1:10 | SRC1 PAS-B | – |
| phenylethylamine-HCl | 1:10, 1:40 | SRC1 PAS-B | – |
| epinephrine | 1:10 | SRC1 PAS-B | – |
| pyrodoxine | 1:10 | SRC1 PAS-B | – |
| lidocaine | 1:10 | SRC1 PAS-B | – |
| nornicotine | 1:10 | SRC1 PAS-B | – |
| carbamazepine | 1:10 | SRC1 PAS-B | – |
| diacetin | 1:10 | SRC1 PAS-B | – |
| all-*trans* retinoic acid | 1:10 | PAS-B-Nurr1 $AF1^{28\text{-}51}$ Fusion | – |
| arachidonic acid | 1:10 | PAS-B-Nurr1 $AF1^{28\text{-}51}$ Fusion | – |

*: all experiments performed at 0.2 mM protein concentration

**: highly reactive compound; all titrations with highly reactive compounds were conducted at lower protein:ligand ratios

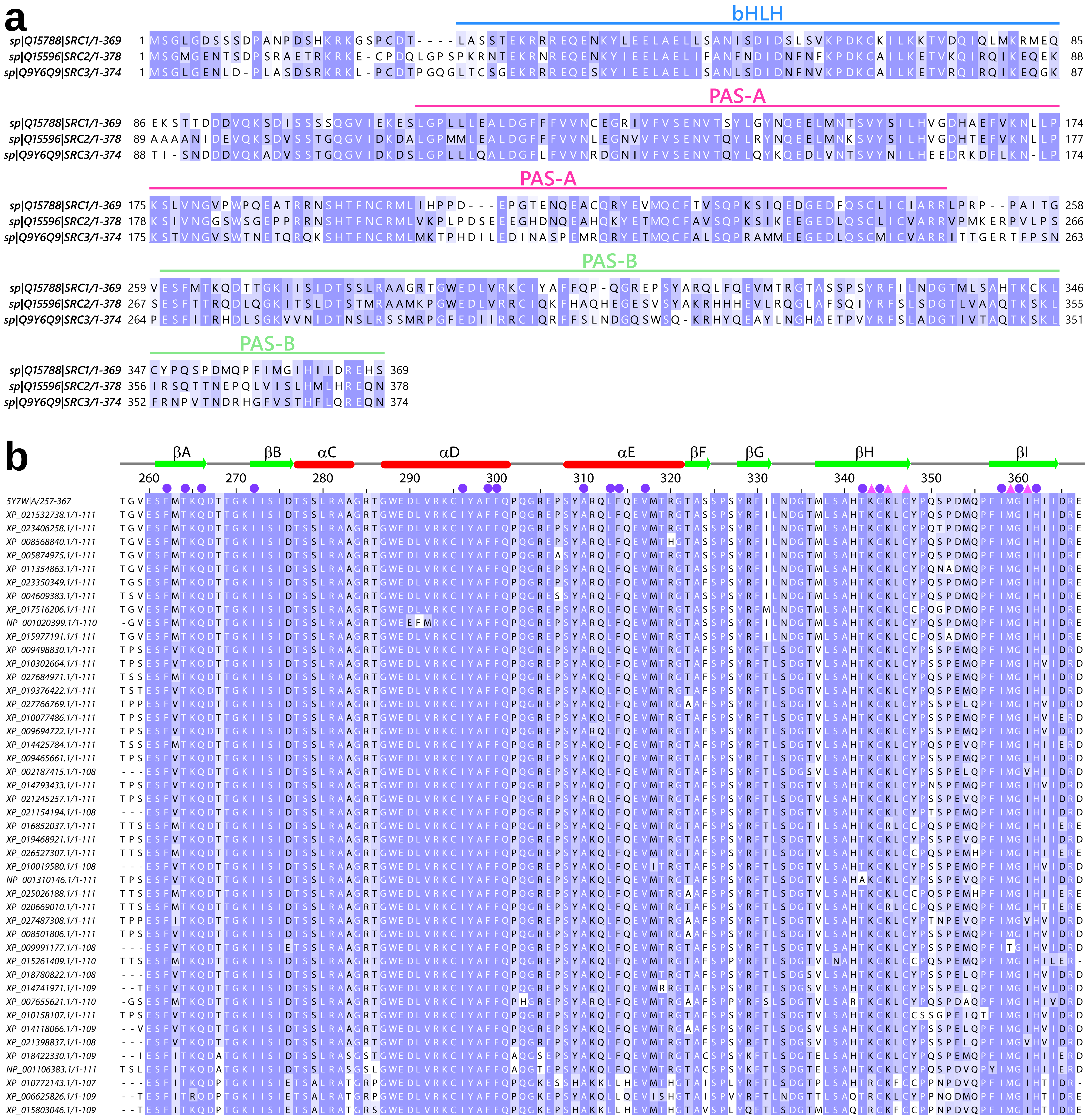


**Supplementary Figure S1**

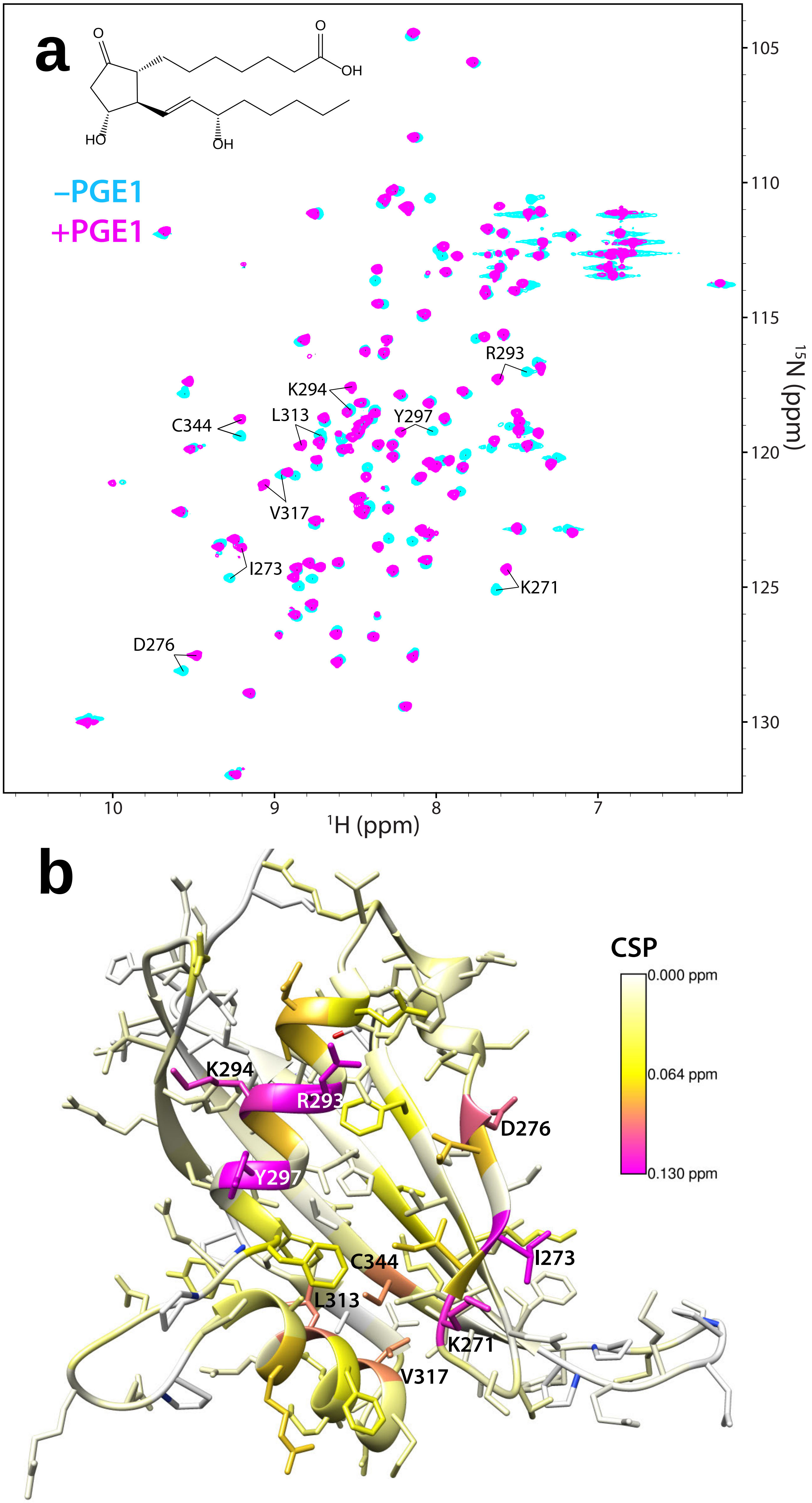


**Supplementary Figure S2**

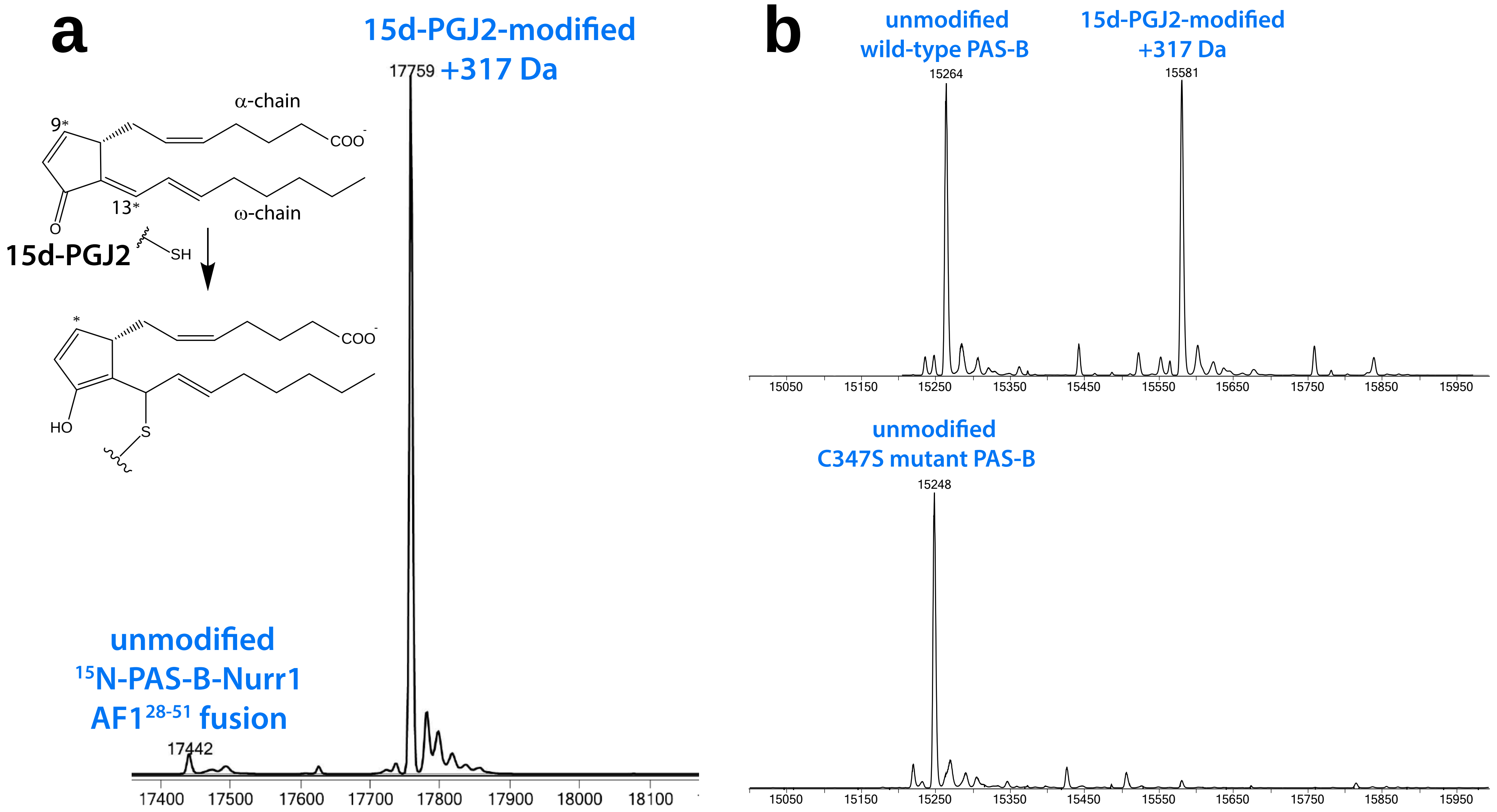


**Supplementary Figure S3**
